# Interpretable Artificial Neural Networks incorporating Bayesian Alphabet Models for Genome-wide Prediction and Association Studies

**DOI:** 10.1101/2021.04.07.438762

**Authors:** Tianjing Zhao, Rohan Fernando, Hao Cheng

## Abstract

In conventional linear models for whole-genome prediction and genome-wide association studies (GWAS), it is usually assumed that the relationship between genotypes and phenotypes is linear. Bayesian neural networks have been used to account for non-linearity such as complex genetic architectures. Here, we introduce a method named NN-Bayes, where “NN” stands for neural networks, and “Bayes” stands for Bayesian Alphabet models, including a collection of Bayesian regression models such as BayesA, BayesB, BayesC, Bayesian LASSO, and BayesR. NN-Bayes incorporates Bayesian Alphabet models into non-linear neural networks via hidden layers between SNPs and observed traits. Thus, NN-Bayes attempts to improve the performance of genome-wide prediction and GWAS by accommodating non-linear relationships between the hidden nodes and the observed trait, while maintaining genomic interpretability through the Bayesian regression models that connect the SNPs to the hidden nodes. For genomic interpretability, the posterior distribution of marker effects in NN-Bayes is inferred by Markov chain Monte Carlo (MCMC) approaches and used for inference of association through posterior inclusion probabilities (PIPs) and window posterior probability of association (WPPA). In simulation studies with dominance and epistatic effects, performance of NN-Bayes was significantly better than conventional linear models for both GWAS and whole-genome prediction, and the differences on prediction accuracy were substantial in magnitude. In real data analyses, for the soy dataset, NN-Bayes achieved significantly higher prediction accuracies than conventional linear models, and results from other four different species showed that NN-Bayes had similar prediction performance to linear models, which is potentially due to the small sample size. Our NN-Bayes is optimized for high-dimensional genomic data and implemented in an open-source package called “JWAS”. NN-Bayes can lead to greater use of Bayesian neural networks to account for non-linear relationships due to its interpretability and computational performance.

## Introduction

Genomic prediction plays an important role in animal and plant breeding by using genomic information to estimate genotypic values or breeding values of complex traits. The adoption of genomic information, such as single-nucleotide polymorphisms (SNPs), has greatly shortened the generation interval and improved the prediction accuracy (Meuwissen *et al.* 2001; Hayes *et al.* 2009a; Heffner *et al.* 2009; Hickey *et al.* 2017). Genome-wide association studies (GWAS) are used to detect associations between SNPs and traits. It has been applied in human diseases (e.g., Ozaki *et al.* (2002); Hirschhorn and Daly (2005); Klein *et al.* (2005); Visscher *et al.* (2012, 2017); Buniello *et al.* (2019)), as well as for traits of interest in animals and plants (e.g., Atwell *et al.* (2010); Korte and Farlow (2013); Sharma *et al.* (2015); Freebern *et al.* (2020)).

Based on the work of Fisher (1918), we typically assume that complex traits are affected by many genes with small additive effects, and that the relationship between genotypes and phenotypes is linear. Based on this assumption, the landmark paper of Meuwissen *et al.* (2001) introduced Bayesian regression models for whole-genome prediction. In Meuwissen *et al.* (2001), they proposed a model referred to as BayesA, where the marker effects were assigned a student-t prior distribution, and a model BayesB, where a-priori only a proportion of markers had nonzero effects. Since Meuwissen *et al.* (2001), several variations to these regression models have been proposed, and, hereafter, they will be collectively referred as Bayesian Alphabet models. Most of the widely-used Bayesian regression models, including BayesA, BayesB, BayesC (Kizilkaya *et al.* 2010; Habier *et al.* 2011), Bayesian LASSO (Park and Casella 2008; Gianola and Fernando 2019), and BayesR (Erbe *et al.* 2012; Moser *et al.* 2015), differ only in the prior used for the marker effects. Another well-known linear model for genomic prediction is genomic best linear unbiased prediction (GBLUP) (Habier *et al.* 2007; VanRaden 2008; Hayes *et al.* 2009b), where a genomic relationship matrix is used to account for the covariances among genetic values. It has been showed that GBLUP is equivalent to a Bayesian regression model with a normal prior for the marker effects (Fernando 1998; Habier *et al.* 2007; Strandén and Garrick 2009). These models, including their variations, have been widely used in GWAS (Fernando *et al.* 2017; Wang *et al.* 2012, 2016; Moser *et al.* 2015; Legarra *et al.* 2018). The assumption of linearity in these analyses, however, may affect their performance on genome-wide prediction and association studies (Nelson *et al.* 2013). Thus, machine learning models have been suggested due to their ability to capture the intricate non-linear relationship between high-dimensional inputs (genotypes) and outputs (phenotypes) (Szymczak *et al.* 2009; Azodi *et al.* 2019).

Artificial neural networks, inspired by information processing of brain, are a subset of machine learning models. Neural networks utilize multi-layer architectures to learn the representation of data (LeCun *et al.* 2015), where the processing layers are composed of neurons (i.e., nodes) converting the input data into a more abstract representation. Neural networks have demonstrated their predictive ability in many fields such as speech recognition, object recognition and detection (LeCun *et al.* 2015). It has also been applied in biological fields such as protein structure prediction, protein classification, and brain decoding (Min *et al.* 2017).

In some genomic prediction studies, neural networks with different architectures were compared to conventional linear models, such as GBLUP and Bayesian regression models. In genomic prediction applications, single hidden layer neural networks are typically applied, but they vary in the number of nodes in the hidden layer (Gianola *et al.* 2011; Okut *et al.* 2011, 2013; Ehret *et al.* 2015). Neural networks achieved higher prediction accuracies in some studies (e.g., Gianola *et al.* (2011); Ma *et al.* (2018)), but a gain in prediction performance was not consistently observed (e.g., Okut *et al.* (2013); Bellot *et al.* (2018); Azodi *et al.* (2019); Abdollahi-Arpanahi *et al.* (2020)), potentially due to the sample sizes in these studies or genetic architecture of the traits (Abdollahi-Arpanahi *et al.* 2020).

Besides the absence of a substantial improvement in prediction accuracy in genomic prediction, a major criticism of neural network models is the lack of interpretability. For example, deep learning (“deep” typically refers to neural networks with more than two hidden layers (Schmidhuber 2015)) is applied in a black-box manner, i.e., it is hard to interpret results biologically for genome-wide analyses, although the neural network architecture may capture the complex relationships between inputs (genotypes) and outputs (phenotypes). Unlike neural networks, conventional linear models provide an explicit interpretation to link phenotypes to genotypes under some assumptions, and thus, they are used in many genome-wide analyses, e.g., GWAS. This lack of interpretability of results from neural networks impedes their application for association studies (Ehret *et al.* 2015; Waldmann 2018). In some studies (e.g., Okut *et al.* (2011); Glória *et al.* (2016); Waldmann (2018)), weights in neural networks were interpreted as regression coefficients to obtain the effect of each SNP, and these effects without significance measures (e.g., *p*-value) were used to rank all SNPs.

Bayesian regularization has been applied to the neural networks to reduce their complexity by treating the networks weights as random variables with prior densities (e.g., Gianola *et al.* (2011); Okut *et al.* (2011, 2013); Glória *et al.* (2016); van Bergen *et al.* (2020)). However, in most Bayesian neural network studies on genomic prediction, in part due to computational reasons, inferences of unknowns were only based on the posterior mode via the maximum a posteriori (MAP) estimator (e.g., Gianola *et al.* (2011); Okut *et al.* (2011, 2013); Glória *et al.* (2016)). Chen *et al.* (2017a) showed that in association studies, the performance of MAP inferences may be inferior to Markov chain Monte Carlo (MCMC) approaches, which exactly estimate the posterior distribution. In Demetci *et al.* (2020), variational inference was used to approximate the posterior distribution of posterior inclusion probabilities (PIPs) for all SNPs. However, simulations in Demetci *et al.* (2020) showed that such approximation may mis-estimate the network weights.

Although Bayesian methods had been proposed for analyses of genetic populations (Dempfle 1977; Gianola and Fernando 1986) in the 1980’s, these methods were not widely adopted until MCMC approaches were introduced to draw inferences from posterior distributions (Wang *et al.* 1994). In the method proposed here, the posterior distribution of marker effects is inferred using MCMC approaches, and it is used to produce significance measures such as posterior inclusion probability (PIP) (Guan and Stephens 2011) and window posterior probability of association (WPPA) (Fernando *et al.* 2017). The method is called NN-Bayes, where “NN” stands for neural networks, and “Bayes” stands for Bayesian Alphabet models, a collection of Bayesian regression models such as BayesA, BayesB, BayesC, Bayesian LASSO, and BayesR. As showed in Figure 1, the framework of NN-Bayes starts with SNPs that are linearly connected to hidden nodes, through a multi-trait, multiple regression model, that are then non-linearly connected to the observed trait. Multi-trait Bayesian Alphabet models (Cheng *et al.* 2018b) will be employed to fit the regression model that connects the SNPs to the hidden nodes. The hidden nodes, sampled via Hamiltonian Monte Carlo, represent unobserved intermediate traits that are nonlinearly connected to the observed trait to accommodate complex genetic architecture including non-additive genetic effects. In summary, NN-Bayes attempts to improve the performance of genome-wide prediction and GWAS by accommodating nonlinear relationships between the hidden nodes and the observed trait, while maintaining genomic interpretability through the Bayesian regression models that connect the SNPs to the hidden nodes.

**Figure 1.**
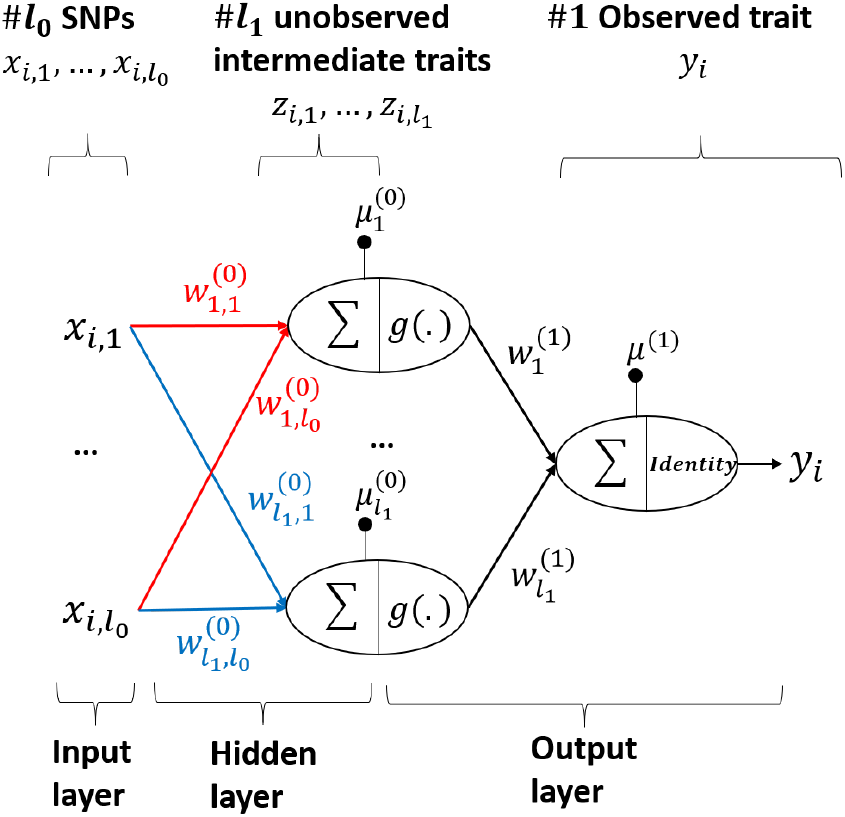
Framework of NN-Bayes. SNPs are linearly connected to unobserved intermediate traits (i.e., hidden nodes). Multitrait Bayesian Alphabet models will be employed to sample marker effects on hidden nodes (i.e., weights between input and hidden layers). Then hidden nodes are non-linearly connected to the observed trait to accommodate complex genetic architecture including non-additive genetic effects, and the non-linearity is achieved by the non-linear function *g*(.).

Compared to other neural network applications of genome-wide prediction and association studies, our NN-Bayes using MCMC approaches is optimized for high-dimensional genomic data to be computationally efficient. In most neural network applications of genomic prediction, neural networks were implemented using existing general purpose software tools which are not optimized for genomic analyses, and it would be time consuming to estimate effects of a large number of genome-wide molecular markers (i.e., weights between input layer and hidden nodes). For example, in van Bergen *et al.* (2020), it took about 4 hours to run 1,000 iterations for a single-layer neural network with 20 hidden nodes using a dataset of 500 individuals and 5,000 SNPs. However, the computing time of our NN-Bayes, implemented in an open-source package “JWAS”(Cheng *et al.* 2018a), is about 2 minutes for such an analysis on a laptop (with 2.6 GHz Intel Core i7 processor), and it scales nearly linearly by the number of observations and the number of markers.

## Materials and Methods

### Bayesian Neural Network Alphabet

Bayesian analyses using NN-Bayes will be demonstrated for a single observed trait. The hierarchical Bayesian model of NN-Bayes is showed in Figure 2. For individual *i*, the observed trait is modeled as

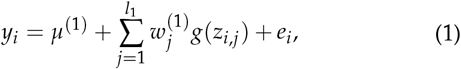

where *y*_*i*_ is the observed phenotype for individual *i*, *μ*^(1)^ is the overall mean, *z*_*i,j*_ is the *jth* hidden node for individual *i*, *g*(.) is an element-wise non-linear function, 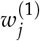 is the effect of *g*(*z*_*i,j*_) on *y*_*i*_, and *e*_*i*_ is a random residual. Non-linearity in the model is achieved through use of the function *g*(.), which is called an activation function in the artificial neural network literature (Leshno *et al.* 1993), and the hyperbolic tangent function was applied in this paper. Flat priors are used for the weights 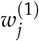 and the overall mean *μ*^(1)^; the prior for *e*_*i*_ is a normal distribution with null mean and unknown variance 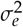, i.e., 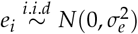. A scaled inverse chi-squared distribution is assigned as the prior for 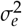, i.e., 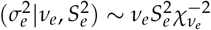.

**Figure 2.**
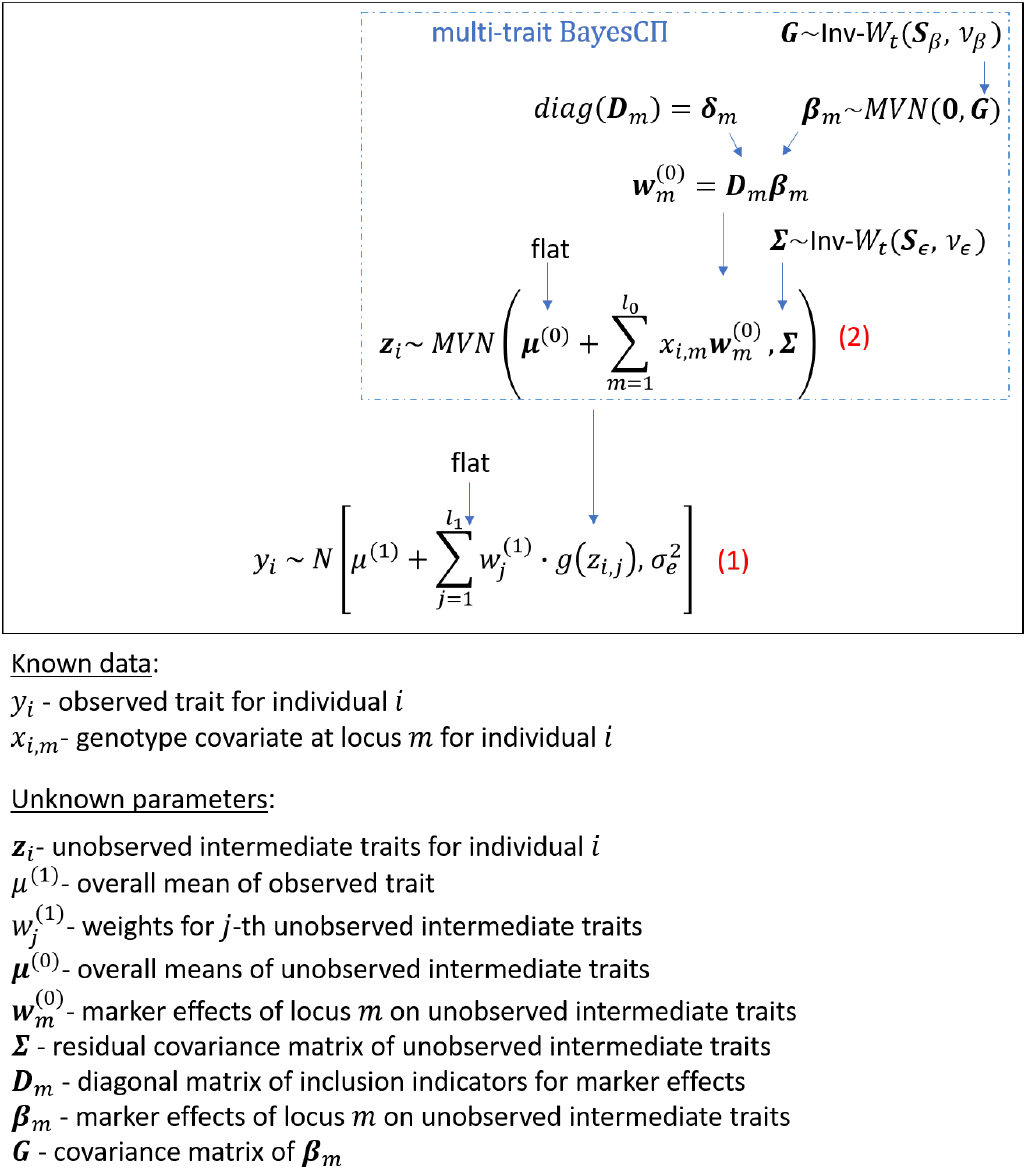
NN-Bayes represented as a hierarchical Bayesian model. Mixture priors in multi-trait BayesCn are used for marker effects on unobserved intermediate traits.

For *i*th individual, the prior for the hidden nodes, 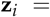 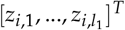, can be presented as a multi-trait Bayesian regression model (Cheng et *al.* 2018b):

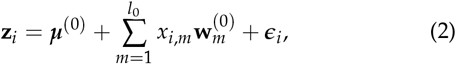

where **z**_**i**_ is the vector of *l*_1_ hidden nodes, which can be thought of as unobserved intermediate traits, for individual *i*, 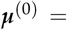 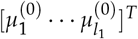 is a vector of overall means for the *l*_1_ hidden nodes, *x*_*i,m*_ is the observed genotype covariate at locus *m* for individual *i* (coded as 0,1,2), 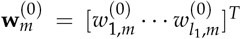 are the marker effects of locus *m* on the *l*_1_ unobserved intermediate traits (i.e.,weights between *m*-th input node and all hidden nodes in the neural network), and *ϵ*_*i*_ is a vector of random residuals for the *l*_1_ hidden nodes.

The overall means, ***μ***^(0)^, are assigned flat priors. Conditional on **Σ**, the residuals, *ϵ*_*i*_, have independently and identically distributed multivariate normal priors with null means and covariance matrix **Σ**, which itself is assumed to have an inverse Wishart prior distribution, 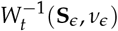. As showed in Cheng *et al.* (2018b), by employing the data augmentation, 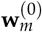 can be written as 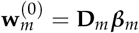, where **D**_*m*_ is a diagonal matrix, where the *k*th diagonal entry indicates whether the marker effects of locus *m* for the unobserved intermediate trait *k* is zero or not. ***β***_*m*_ follows some multivariate distribution, e.g, the multivariate normal distribution in multi-trait BayesCΠ or the multivariate t distribution in multi-trait BayesB. Note that multi-trait Bayesian regression models may be computationally intensive if a large number of hidden nodes are used. In this case, we assume that hidden nodes are independent such that multiple single-trait Bayesian regression models can be used in parallel at each iteration.

In Figure 1, we also present the hierarchical Bayesian model described above as a single hidden layer feed-forward Bayesian neural network. This architecture is typical for neural networks used in genomic prediction (Gianola et *al.* 2011; Okut *et al.* 2011, 2013; Ehret *et al.* 2015).

In the neural network, for each observation, the input layer is composed of a vector of *l*_0_ molecular markers (coded as 0, 1, 2), and the output layer is the observed trait. Nodes in the hidden layer, between input and output layer, represent unobserved intermediate traits, and the relationship between hidden nodes and observed trait is non-linear. The weights between the input layer and the hidden layer represent marker effects on unobserved intermediate traits, the “biases” of hidden nodes can be regarded as the overall means. The weights and bias between the hidden layer and the output layer help to define the non-linear relationship between unobserved intermediate traits and observed trait. Note that extra input nodes can be included in the neural network if additional random or fixed effects are fitted in the model.

### Inference using Markov chain Monte Carlo

From a Bayesian perspective, inferences of unknowns are based on their posterior distributions. However, closed-form expressions of those posteriors are usually unavailable. Thus, in practice, samples are obtained using MCMC techniques, where statistics computed from the resulting Markov chain converge to those from the posterior as chain length increases (Norris 1998; Sorensen and Gianola 2007). Here, to construct the Markov chain, we will use Gibbs sampling, where each unknown variable or block of variables is sampled from its full conditional distribution, conditioned on the observed data and the latest samples of all the other unknowns.

Given the sampled values of hidden nodes **z**_*i*_ in equation (2), unknown parameters between the input layer and the hidden layer do not depend on the observed trait. Sampling of those unknowns, including the overall means ***μ***^(0)^, the marker effects 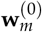, and the residual covariance matrix **Σ**, given the sampled values of **z**_*i*_, is based on the multi-trait Bayesian regression models described in Cheng *et al.* (2018b). Given the sampled values for the hidden nodes (after activation) *g*(**z**_*i*_) and output node *y*_*i*_, the sampling of weights and bias between the hidden layer and the output layer is straightforward (derivations are given in the Appendix). As showed below, we consider Hamiltonian Monte Carlo to draw samples for the hidden nodes. The sampler is implemented in the open-source package “JWAS” (Cheng *et al.* 2018a).

### Sampling hidden nodes using Hamiltonian Monte Carlo

Hamiltonian Monte Carlo (HMC) is used to sample hidden nodes, i.e., **z**_*i*_. In HMC, each unknown parameter is paired with a “momentum” variable ***ϕ***. The HMC constructs the Markov chain by a series of iterations. A useful introduction to the principles and concepts underlying HMC is given by Betancourt (2017). Following notations in Gelman et al. (2013), there are three steps in each iteration of the HMC:

1. updating the momentum variable independently of the current values of the paired parameter, i.e., ***ϕ*** ~ *MVN*(0, **M**).
2. updating (**z**_*i*_, ***ϕ***_*i*_) via *L* “leapfrog steps”. In each leapfrog step, **z**_*i*_ and ***ϕ***_*i*_ are updated dependently and scaled by *ε*. The leapfrog step below is repeated *L* times:

a. 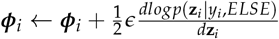
b. ***z***_*i*_←**z**_*i*_ + *ϵ***M**^−1^*ϕ_i_*;
c. 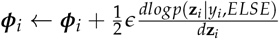. The resulting state at the end of L repetitions will be denoted as 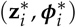.
3. calculating the acceptance rate, *r*, and above resulting state will be accepted with probability *min* (1, *r*).

In our method, the gradient of log full conditional posterior distribution of **z**_*i*_ is:

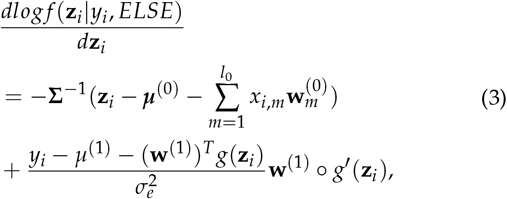

where 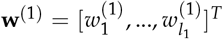 is the vector of weights between hidden nodes and observed trait, *ELSE* denotes the sampled values of remaining unknowns, and the symbol “ ο ” denotes the element-wise production.

In the round t of HMC, the acceptance rate, *r* is:

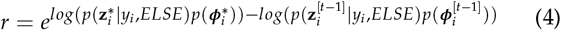

A detailed derivation can be found in the Appendix. In our analyses, 10 leapfrog steps was applied in each iteration of HMC, i.e., *L* = 10, the scale parameter *ϵ* was 0.1, and **M** was set as an identity matrix.

### NN-Bayes for Genomic Prediction

The MCMC sample from the posterior distribution of genotypic values for individuals of interest is obtained as

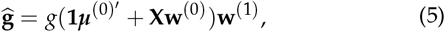

where **X** is the genotype covariate matrix for individuals of interest, **w**^(0)^ is a matrix of size *p*-by-*l*_1_ of samples of marker effects on the *l*_1_ hidden nodes, ***μ***^(0)^ is a vector of samples of overall means for the *l*_1_ hidden nodes, and **w**^(1)^ is a vector of samples of *l*_1_ weights between hidden nodes and the observed trait, and *g*(.) is a non-linear function. The prediction accuracy is calculated as the Pearson correlation between the posterior mean of genotypic values and the phenotypic values in the validation dataset.

### NN-Bayes for Genome-wide Association Studies

In NN-Bayes, the PIP for each single marker is computed as the frequency that its effect is non-zero on at least one of hidden nodes. Here we prefer the use of the posterior probability of association of a genomic window (WPPA) (Fernando *et al.* 2017) to account for the fact that highly correlated SNPs within a genomic window jointly affect the phenotype, and it is difficult to identify the effect of a single marker (Hayes et al. 2010).

Given that an investigator is interested in identifying genomic segments that explain more than a proportion T of the genetic variance, WPPA is defined as the posterior probability of this event (Fernando *et al.* 2017). To estimate WPPA, first, MCMC samples of genotypic values for all individuals are obtained from their posterior distribution, using equation (5). Next, using the notation in equation (5), the genotypic values that are attributed to genomic window *t* are sampled from their posterior distribution as

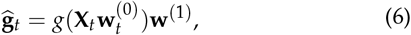

where **X**_*t*_ is the genotype covariate matrix of markers in window *t*, and 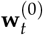 represents the samples of marker effects in window *t* on hidden nodes. The proportion of the genetic variance explained by the genomic window *t*, *q*_*t*_, is now sampled as 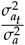, where 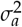 and 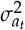 are the variances of a random sample from **g** and **g**_*t*_, respectively. Then, the posterior probability that window *t* accounts for more than a proportion *T* of the genetic variance (i.e., WPPA) can be estimated from the MCMC samples by counting the number of samples where *q*_*t*_ > *T* (Fernando and Garrick 2013; Chen et al. 2017b; Lloyd-Jones *et al.* 2017). In this paper, non-overlapping windows of 1 Mb were used to identify genomic windows that explain over 1% of the total genetic variance (i.e., *T*=0.01)

### Data Analysis

#### Simulated data

Real pig genotypes in Duarte et al. (2014) were used to simulate phenotypes with dominance and epistatic effects. All SNP markers on the chromosome 1 were used, resulting a genotypic data with 5,023 markers for 928 individuals. A random sample of five percent of these 5,023 markers were selected as quantitative trait loci (QTL). Following the simulation in van Bergen et al. (2020), the additive effects (*a*) of QTL were sampled from a univariate normal distribution with null mean and variance one. Dominance factors (*δ*) were sampled from a univariate normal distribution with mean 1.2 and standard deviation 0.3. Dominance effects (*d*) were computed as *δ*|a|. The epistatic factors (*γ*) of all pairwise combinations of QTL were sampled from a univariate normal distribution with null mean and variance one. The epistatic effect (*ϵ*) between *i*th and *j*th QTL was computed as 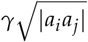. The complementary epistasis scenario in van Bergen *et al.* (2020) was used to simulate the total genetic value. Phenotypes were simulated with a broad-sense heritability of 0.5 and were scaled to have a phenotypic variance equal to one. Based on the simulation processes described above, 20 different datasets were simulated.

#### Real data

Publicly available genotypic and phenotypic data of multiple species were used to compare NN-Bayes and conventional linear models. These datasets include the pig dataset from Duarte et al. (2014), and data for soy, maize, switchgrass, and spruce from Azodi et al. (2019). Traits used in our analyses were the 13 week tenth rib backfat (BF), yield (YLD), wood density (DE) and flowering time (FT). The description of each dataset, such as the number of SNP markers and the number of observations, are showed in Table 1.

**Table 1.**
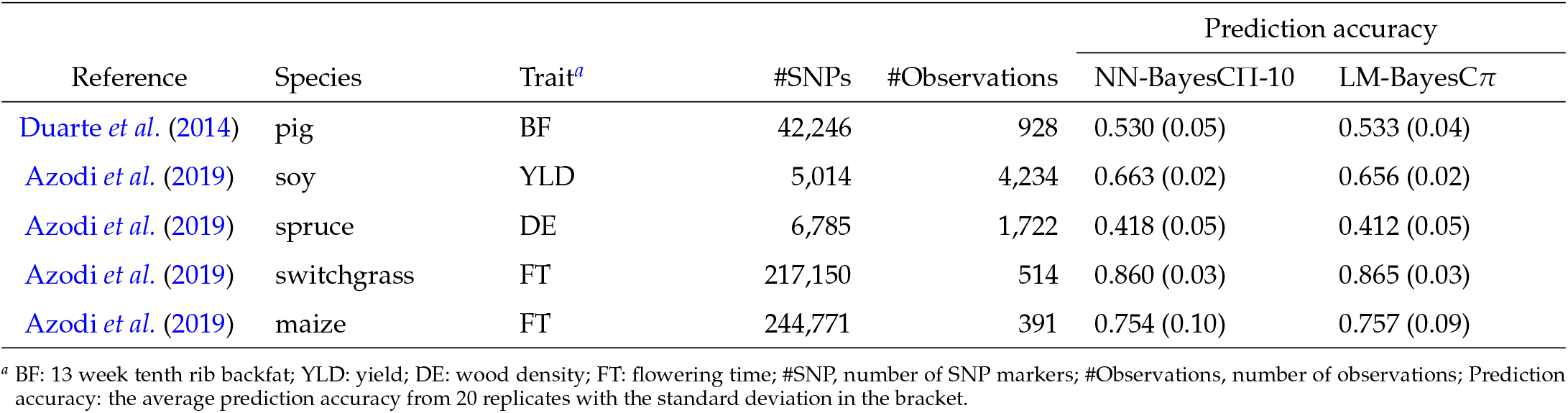
Comparison among five species in prediction accuracies of NN-Bayes composed of 10 hidden nodes with BayesCΠ prior (NN-BayesCΠ-10) and a linear model with BayesC*π* prior (LM-BayesCπ).

#### Genomic Prediction

20 and 50 replicates were applied for each real and simulated dataset, respectively. In each replicate, a random subset of 80% of all observations were used for training, and the remaining were used for validation. The prediction accuracy is calculated as the Pearson correlation between the posterior mean of genotypic values and the phenotypic values in the validation dataset. In NN-Bayes, different priors for marker effects (i.e., weights between inputs and hidden nodes) in Bayesian Alphabet models (i.e., RR-BLUP, BayesA, BayesB, BayesC*π*, and Bayesian LASSO) were used. Linear models with priors in Bayesian Alphabet models were also performed. Different number of hidden nodes (i.e., 2, 3, 5 and 10) were tested in the analyses. Chains of length 10,000 and 20,000 were applied to simulated datasets and real datasets, respectively, to ensure the convergence.

#### Genome-wide association studies

For each dataset, PIP and WPPA were used for association inference. In the simulated study, non-overlapping windows of 1 Mb were tested. Genomic windows that explain over 1% of the total genetic variance were assumed to be of potential interest (i.e., *T*=0.01). The area under the receiver operating characteristic curve (AUC) was calculated using the R package pROC (Robin et al. 2011) to assess the performance of NN-Bayes in GWAS.

#### Data availability

The genotypic and phenotypic data used in the real data analyses are publicly available in Duarte et al. (2014) and Azodi et al. (2019). The authors state that all data necessary for confirming the conclusions presented in the article are represented fully within the article.

## Results

### Genomic Prediction

#### Simulated data

Overall, the prediction accuracies of NN-Bayes were significantly higher than those of conventional linear models, and the differences were substantial in magnitude. The number of hidden nodes did not significantly affect the performance of NN-Bayes. No significant differences were found with different priors for marker effects used in NN-Bayes. Here we only present the comparison of prediction accuracy between NN-Bayes composed of 10 hidden nodes with BayesCΠ priors (named as NN-BayesCΠ-10) and the linear model with BayesC*π* prior (LM-BayesCπ). In Figure 3, the 20 simulated datasets were distinguished by color. Prediction accuracies of NN-BayesCΠ-10 were higher than LM-BayesC*π* in 949 out of 1,000 validation sets (i.e., above the diagonal black line), and the prediction accuracy for NN-BayesCΠ-10 was significantly higher than for LM-BayesC*π* in 16 out of 20 simulated datasets under the t-test with a significance level of 0.05. The results show that our NN-Bayes has the potential to improve the prediction accuracy for a trait with non-additive genetic effects.

**Figure 3.**
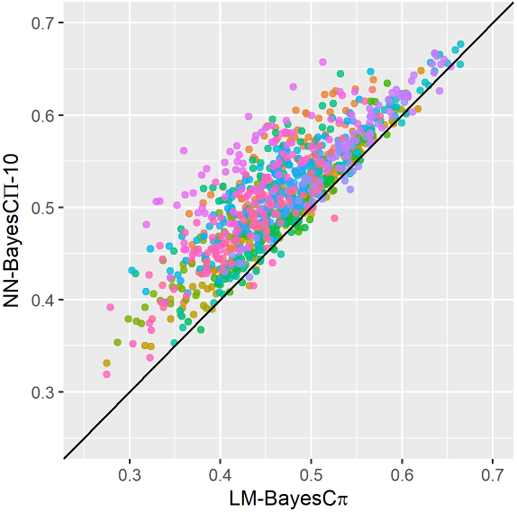
The prediction accuracy of NN-Bayes composed of 10 hidden nodes with BayesCΠ priors (NN-BayesCΠ-10) versus a linear model with BayesC*π* prior (LM-BayesCπ) on scenarios with both additive and non-additive effects (i.e., dominance and epistasis). 20 simulated datasets were distinguished by color. For each dataset, 50 replicates for validation were applied. The diagonal line is used for reference such that a dot above the line represents a validation with higher accuracy for NN-Bayes.

#### Real data

In each of the five species, the prediction accuracies of NN-Bayes with different numbers of hidden nodes were not significantly different under the t-test with a significance level of 0. 05. Significant differences were found among different priors for marker effects in the pig data. Comparison of prediction accuracies among all methods is showed in Table 2 in the Appendix. In detail, NN-Bayes using BayesA and BayesB priors performed better than methods using other priors in the pig data. The prediction accuracies of NN-Bayes composed of 10 hidden nodes with BayesCΠ priors (named as NN-BayesCΠ-10) and the linear model with BayesC*π* prior (LM-BayesCπ) are showed in the last column of Table 1. Among the five species, the prediction accuracies for NN-BayesCΠ-10 were higher than those for LM-BayesC*π* when the ratio between number of observations and number of markers was relatively high (e.g., soy and spruce). However, the prediction accuracies of NN-BayesCΠ-10 were not significantly different from those for LM-BayesC*π* under the t-test with a significance level of 0.05. This may be because the sample size is not large enough for accurate parameter estimates, or the architecture of the neural network may not be sufficiently complex to represent the intricate hidden relationships between genotypes and phenotype. Alternatively, the additive genetic effects may have already accounted for a majority proportion of the variation, so the improvement based on non-additive effects is limited. When we considered a significance level of 0.1, the prediction accuracies for NN-Bayes composed of 10 hidden nodes with priors for marker effects in BayesCΠ or RR-BLUP were significantly higher than conventional linear models for the soy dataset. As showed in Figure 4, for the soy dataset, in 15 out of 20 validation sets, the NN-BayesCΠ-10 had a higher prediction accuracy than the LM-BayesC*π* (i.e., above the diagonal black line).

**Table 2.**
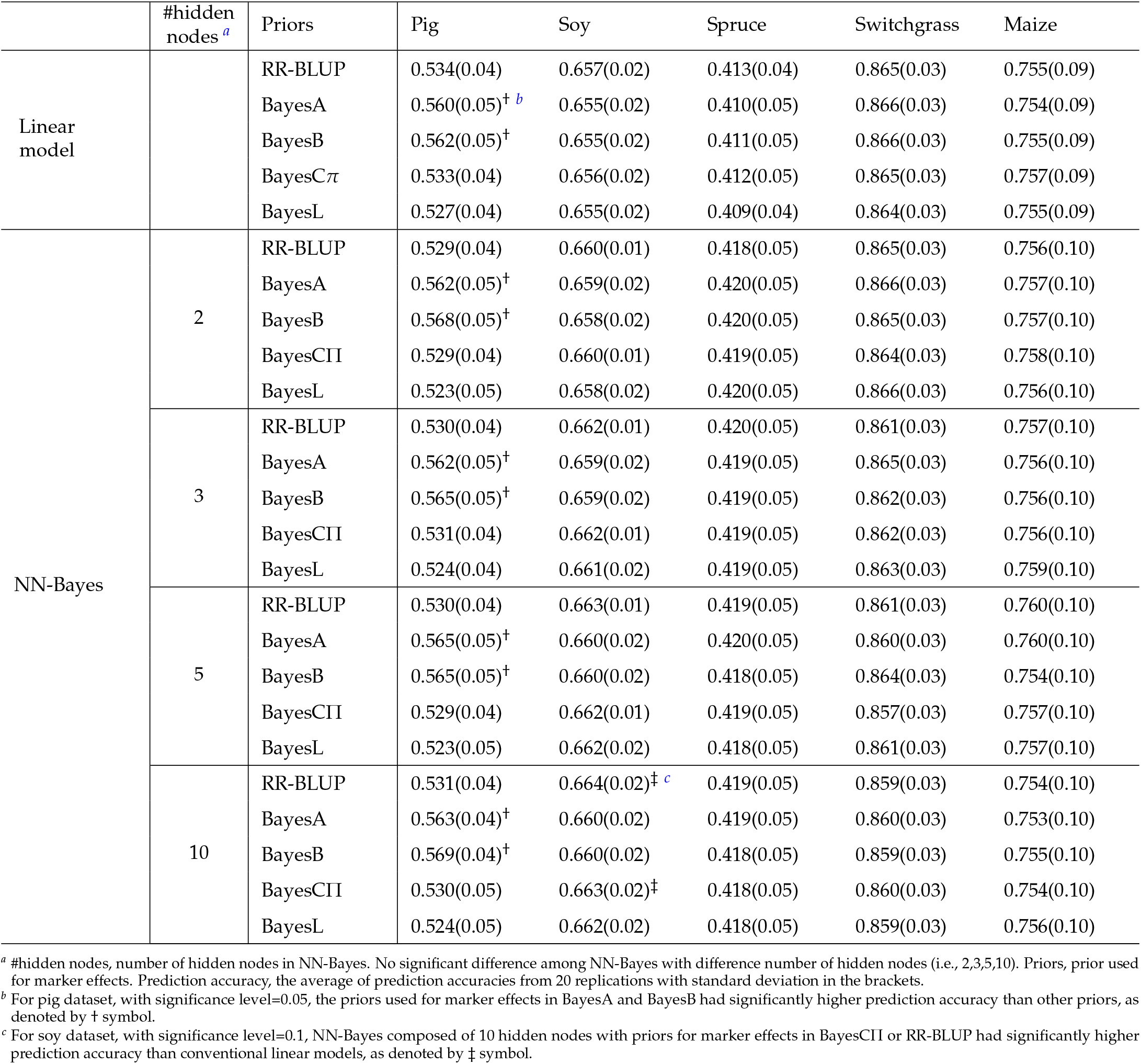
Prediction accuracy of NN-Bayes and linear models for real datasets.

**Figure 4.**
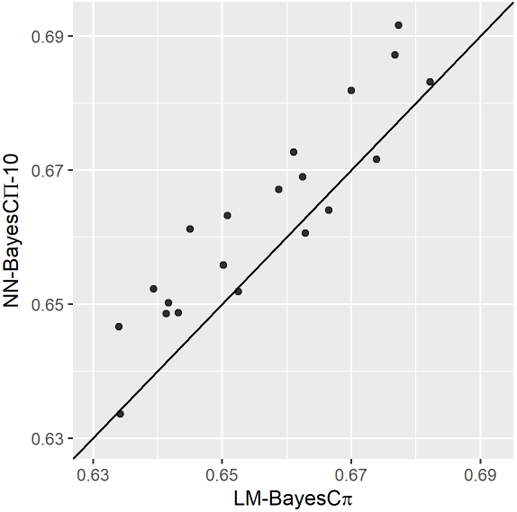
The prediction accuracy of NN-Bayes composed of 10 hidden nodes with BayesCΠ priors (NN-BayesCΠ-10) versus the linear model with BayesC*π* prior (LM-BayesCπ) for the soy dataset. 20 replicates for validation were applied. The diagonal line is used for reference such that a dot above the line represents a validation with higher accuracy for NN-Bayes.

### Genome-wide association studies

#### Simulated Data

GWAS was conducted on 20 simulated datasets. NN-Bayes composed of 2 hidden nodes with priors for marker effects in BayesCΠ (named as NN-BayesCΠ-2) was tested and compared to results from a linear model with BayesC*π* prior (LM-BayesCπ). Chain of length 500,000 with 200,000 burn-in was used to ensure the convergence, and samples were saved for every 50 iterations.

For both NN-BayesCΠ-2 and LM-BayesCπ, when PIP was used for association inference, the AUC was around 0.5, indicating the performance of PIP was similar to a random classifier. Thus, we will only present the GWAS results using WPPA in all analyses. The AUC of NN-BayesCΠ-2 was significantly higher than that of LM-BayesCπ, under the t-test with a significance level of 0.15. As showed in Figure 5, the AUC for NN-BayesCΠ-2 was higher than that for LM-BayesC*π* in 16 out of the 20 simulated datasets (i.e., above the diagonal black line).

**Figure 5.**
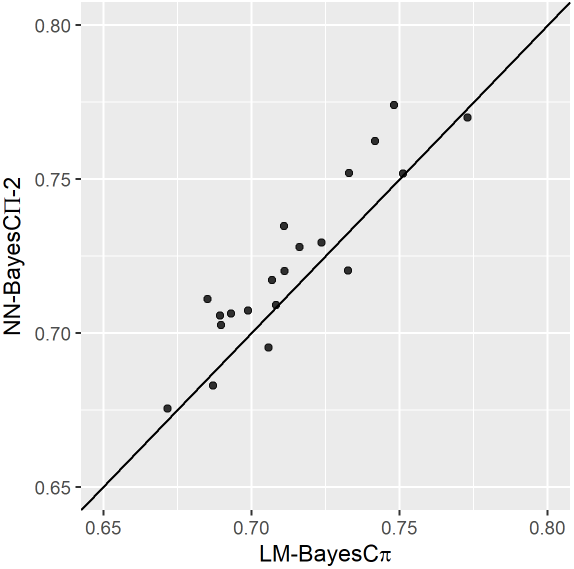
The AUC of GWAS results on 20 simulated datasets for NN-Bayes composed of 2 hidden nodes with BayesCΠ priors (NN-BayesCΠ-2) versus a linear model with BayesC*π* prior (LM-BayesCπ), on scenarios with both additive and non-additive effects (i.e., dominance and epistasis). Inference of association is based on genomic windows. The diagonal line is used for reference such that a dot above the line represents a simulated dateset with higher AUC for NN-Bayes.

#### Real data

GWAS was conducted on the real pig dataset. NN-Bayes composed of 2 hidden nodes with priors for marker effects in BayesCΠ (named as NN-BayesCΠ-2) was tested. Chain of length 1 million with 500,000 burn-in was used to ensure the convergence, and samples were saved for every 10 iterations.

For NN-BayesCΠ-2, a WPPA > 0.95 was used to identify significant associations because it results in controlling the proportion of false positives to ≤ 0.05 (Fernando et al. 2017). The GWAS results from NN-BayesCΠ-2 had a similar pattern as those in Duarte et al. (2014). The significant window on chromosome 6 was identified, which is consistent with the GWAS results in Duarte et al. (2014).

## Discussion

In conventional linear models used for genomic prediction, it is usually assumed that complex traits are affected by many genes with small additive effects, and that the relationship between genotypes and phenotypes is linear. To account for non-linear relationships such as non-additive genetic effects, we proposed a method named NN-Bayes, where “NN” stands for neural networks, and “Bayes” stands for Bayesian Alphabet models, a collection of Bayesian regression models such as BayesA, BayesB, BayesC, BayesR, and Bayesian LASSO. NN-Bayes incorporates Bayesian Alphabet models into non-linear neural networks via hidden layers between SNPs and observed traits. Priors in multitrait Bayesian Alphabet models are assumed for marker effects on hidden nodes, and flexible non-linear relationships between hidden nodes and the observed trait are assumed through activation functions, e.g., the hyperbolic tangent function. Thus, NN-Bayes attempts to improve the accuracy of prediction by accommodating non-linear relationships between the hidden nodes and the observed trait, while maintaining genomic interpretability through the Bayesian regression methods that connect the SNPs to the hidden nodes. Compared to other neural network applications on genomic prediction, NN-Bayes has been optimized for MCMC based inference with high-dimensional genomic data, and it is thus more computationally efficient. Our NN-Bayes is implemented in an open-source package called “JWAS” (Cheng *et al.* 2018a).

In the analyses of the simulated data, where the phenotypes were simulated with both additive and non-additive effects (i.e., dominance and epistasis), performance of NN-Bayes was significantly better than conventional linear models for both GWAS and whole-genome prediction, and the differences on prediction accuracy were substantial in magnitude. In real data analyses, for the soy dataset, NN-Bayes was able to achieve significantly higher prediction accuracies than conventional linear models, and results from other four different species showed that NN-Bayes had similar prediction performance to linear models. Results in Azodi *et al.* (2019) showed that artificial neural networks did not tend to outperform linear models for the multiple plant datasets used in our analyses. This may be because the sample size is not large enough for accurate parameter estimates, or the architecture of the neural network may not be sufficiently complex to represent the intricate hidden relationships between genotypes and phenotype. Alternatively, the additive genetic effects may have already accounted for a majority proportion of the variation, so the improvement based on non-additive effects is limited.

In most neural network applications on genomic prediction, single hidden layer, feed-forward neural networks have been used (e.g., Gianola *et al.* (2011); Okut *et al.* (2011,2013); Ehret *et al.* (2015)). These neural networks are often trained by gradient-based back-propagation, and the training process stops when it reaches the maximum number of training iterations or stops early when a criterion is met, for example, the optimal mean squared error (MSE) on a validation dataset. To reduce the overfitting in such neural networks with large numbers of molecular markers as input, regularization technology, such as Bayesian regularization (Gianola *et al.* 2011) and *L*_1_-norm penalty on unknown parameters (Wang *et al.* 2018), are usually applied. In Bayesian regularized neural networks, the effective number of parameters were similar when different numbers of neurons were used in the middle layer (Gianola *et al.* 2011; Okut *et al.* 2013; Ehret *et al.* 2015) and were usually much smaller than the sample size. Moreover, dropout may be applied to reduce the complexity of neural networks by setting a random proportion of weights to zero in each training iteration (Waldmann 2018; Azodi *et al.* 2019). In van Bergen *et al.* (2020), variable selection was used in a Bayesian neural network by fitting a “marker selection vector” into the Bayesian neural networks.

Most Bayesian neural network studies on genomic prediction heavily rely on approximations in part because they were performed using general algorithms implemented in existing software tools that were not optimized for genomic data analyses. For example, in Gianola *et al.* (2011) and Okut *et al.* (2013), the rationale of using relationship matrix as input of neural networks is from the representation of the infinitesimal model as a regression on relationship matrix, so weights connecting the input and hidden layers should follow the prior distribution of 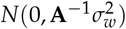, where **A** is the relationship matrix. However, due to the use of MATLAB (Demuth and Beale 2009) in Gianola *et al.* (2011) and Okut *et al.* (2013), this prior was restricted in the form of 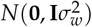. In addition to approximations, it is usually computationally intensive or infeasible to analyze large datasets on genomic prediction using general algorithms implemented in existing software tools. Okut *et al.* (2013) reported that MATLAB required about 2 hours to train a neural network with ~ 2500 SNPs as input, and van Bergen *et al.* (2020) reported that PyMC3 (Salvatier *et al.* 2016) took about 4 hours with 500 individuals and 5000 SNPs. For a dense genotype, such as the Jersey cow data with 35,798 SNPs, the relationship matrix of size 297 was used as input to make computations feasible in MATLAB (Gianola *et al.* 2011). These methods (Gianola *et al.* 2011; Okut *et al.* 2011, 2013; Ehret *et al.* 2015; Bellot *et al.* 2018; Azodi *et al.* 2019) were not used for GWAS due to the lack of interpretability. Our NN-Bayes using MCMC approaches is computationally efficient and can be further sped up by a recently developed parallel computing strategy (Zhao *et al.* 2020).

## Appendix MCMC in NN-Bayes

## Sampling weights between hidden layers and output layers

The full conditional posterior distribution of *μ*^(1)^ and w^(1)^, 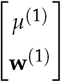, is a multivariate normal distribution with mean:

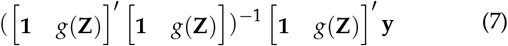

and covariance matrix:

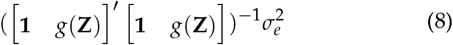

where **y** is a vector of *n* phenotypes for the observed trait, and **Z** is a matrix of size *n*-by-*l*_1_ for hidden nodes.

## Sampling hidden nodes using Hamiltonian Monte Carlo

In the round *t* of HMC, the acceptance rate, *r*, can be expressed as:

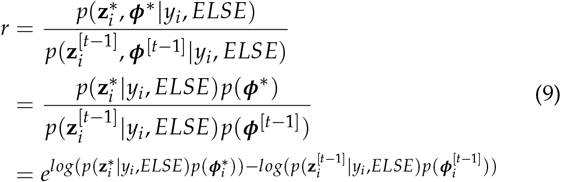

As showed in equation (3) and (9), the log full conditional posterior distribution of the hidden nodes and their gradients are required in HMC. Following equation (3), the full conditional posterior distribution of the hidden nodes for individual *i*, i.e., z_*i*_, can be expressed as:

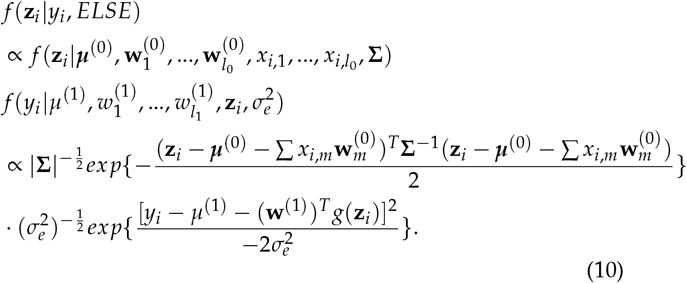

Thus, the log full conditional posterior distribution of z_*i*_ is:

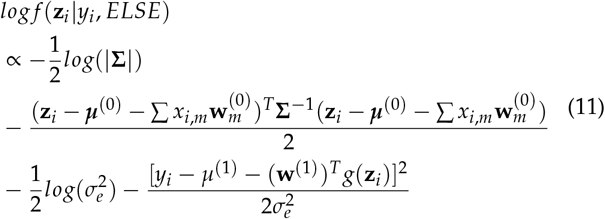

Thus, the gradient of the log-full conditional posterior distribution
of z_*i*_ is:

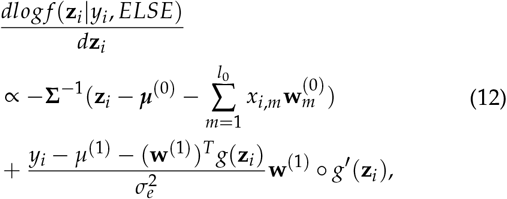

where the symbol “ o “ denotes the element-wise production.

## Sampling residual variance of the observed trait

The full conditional posterior distribution of 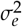 is a scaled inverse chi-squared distribution with *n* + *v*_*e*_ degrees of freedom and scale parameter 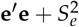, where the **e** is **y** - **1***μ*^(1)^ - g(**Z**)**w**^(1)^.

## Genomic Prediction using real datasets

Prediction accuracies for each real dataset are showed in table 2. In each of the five species, the prediction accuracies of NN-Bayes with different numbers of hidden nodes were not significantly different under the t-test with a significance level of 0.05. Significant differences were found among different priors for marker effects in the pig data.

